# Spatiotemporal representations of contextual associations for real-world objects

**DOI:** 10.1101/2025.10.20.683392

**Authors:** Lu-Chun Yeh, Marius V. Peelen, Daniel Kaiser

## Abstract

In the real world, objects always appear in context. Many objects are reliably associated with a certain scene context (e.g., pots appear in kitchens) and other objects that appear in the same context (e.g., pans appear together with pots). Previous neuroimaging work suggests that such contextual associations shape the neural representation of isolated objects even in the absence of the scene context. Yet, three key questions remain unanswered: (1) How do representations of contextual associations relate to perceptual and categorical representation in visual cortex, (2) how do they emerge across time, and (3) how are they mechanistically implemented? To answer these questions, we recorded fMRI and EEG while participants (human, both sexes) viewed isolated objects stemming from two scene contexts. Multivariate pattern analysis on the neural data revealed that objects from the same context were coded more similarly than objects from different contexts in object-selective LOC and scene-selective PPA, even when systematically controlling for perceptual and categorical similarities. Such contextual relation representations emerged relatively late during visual processing (i.e., after perceptual and categorical representations), specifically in the anterior PPA, and likely through a mixture of object-to-object and object-to-scene associations. Together, our results demonstrate that contextual relation representations emerged for isolated objects, and without a task that encourages their formation, suggesting that objects automatically activate context frames that support visual cognition in real-world environments.

## Introduction

Many real-world objects are specifically associated with the scenes they appear in, creating contextual associations both between objects and scenes and between objects (Bar, 2004; Oliva & Torralba, 2007). For example, a rolling pin is typically found in a kitchen, among other kitchen utensils or food. Previous research suggests that contextual associations are activated spontaneously when viewing objects and scenes (Biederman, 1972; Cornelissen & Võ, 2017; Nah et al, 2021), supported by “context frames” that store the relevant associations in memory (Bar, 2004).

Neuroimaging studies showed that contextual associations shape the cortical representation of individual objects in isolation. Bar and Aminoff (2003) reported that objects with stronger contextual associations (i.e., cooking pots implying a kitchen context) elicited greater activation in the scene-selective parahippocampal place area (PPA) and object-selective area lateral occipital cortex (LOC) than objects without such associations (i.e., bins found across a range of scene contexts). More recently, Bonner and Epstein (2021) demonstrated that the PPA represents object co-occurrence statistics, such that objects sharing contextual associations (e.g., an oven and a dishwasher, typically found together in a kitchen) elicit more similar neural responses than those not sharing such associations (e.g., an oven and an armchair, typically found in different scene contexts). This finding suggests that object representations in scene-selective cortex are organized by contextual associations—likely providing a representational link between scene and object processing. However, three key questions about representations of contextual associations remain unanswered. These pertain to (1) the relation of contextual representations to perceptual and categorical representations, (2) the temporal emergence of contextual representations, and (3) the neural mechanisms that give rise to contextual representations.

First, the exact relationship between contextual relation representations and other representational organizations in visual cortex, such as perceptual and categorical representations, remains unclear. Disentangling these representations is critical, because objects that frequently co-occur in the same context often share perceptual and categorical similarity. For example, most cooking utensils are metallic and shiny tools, conflating perceptual, categorical, and contextual similarities. Without comprehensively controlling the potentially confounding attributes, it remains unclear whether representational similarities truly reflect the objects’ associated context.

Second, it is unclear how contextual relation representations emerge across time. They could emerge relatively late during visual processing, after the computation of perceptual and categorical object features. Such late contextual representations might be driven by more abstract and memory-related processes (Bar & Aminoff, 2003, Bar et al., 2008; Aminoff et al., 2013) that are coded in anterior subregions of the PPA (Baldassano et al., 2013, 2016). Alternatively, contextual association may emerge concurrently with categorical representations, suggesting parallel computations of object category and context in object- and scene-selective cortex.

Third, different mechanisms could mediate the representation of contextual associations. The representations could emerge from a direct co-activation of object representations, akin to a spread of activation across objects from the same context (Bonner & Epstein, 2021; Nah & Geng, 2022), or form shared activation of scene-category representations, consistent with PPA’s role in scene categorization (Epstein & Kanwisher, 1998; Walther et al., 2009). Notably, these two mechanisms are complementary and could both contribute at different processing stages.

We addressed these three open issues by showing participants individual object images from a stimulus set that orthogonally manipulated contextual and categorical relationships while tightly controlling for perceptual similarity. Using multivariate pattern analysis (MVPA) on fMRI and EEG data, we mapped neural representations of contextual associations across space and time. Our results provide three key insights: First, contextual associations are encoded in object- and scene-selective areas, when systematically controlling for perceptual and categorical similarities. Second, such contextual relation representations emerge relatively late in visual processing, after perceptual and categorical object processing, and more strongly in anterior PPA. Third, while initial representations of contextual associations are likely mediated by a co-activation of contextually related objects, later representations may also reflect a co-activation of associated scene category representations.

## Materials And Methods

### Participants

Thirty-four healthy volunteers (22 females, 11 males, and one preferred not to reveal, age =25.21 ± 4.44 years) participated in the EEG experiment, and another 33 (19 females, age =24.14 ± 3.34 years) participated in the fMRI experiment. Sample sizes were chosen to obtain ~80% power for detecting a hypothetical medium effect of d = 0.5 at p < 0.05 (two-sided t-test). All participants were native German speakers and had normal or corrected-to-normal vision. They provided written informed consent prior to participation and were compensated at a rate of 10 euros per hour. The study was approved by the Ethics Committee of Justus Liebig University Giessen and conducted in accordance with the 6th revision of the Declaration of Helsinki. One fMRI participant did not complete all experimental runs and was therefore excluded from the analyses.

### Stimuli

A total of 24 real-world objects were selected, stemming from two contexts (12 kitchen objects and 12 garden objects) and two categories (12 tools and 12 non-tools). These objects were grouped into six sets of four items each, each of which contained one object per condition (i.e., 1 kitchen tool, 1 garden tool, 1 kitchen non-tool, and 1 garden non-tool). Within each set, objects were matched for their overall shape (see Fig. 1A). There were four exemplars per object, yielding a total of 96 unique stimuli. All stimuli were positioned against a white square background (5° × 5° visual angle). The resulting images were gray-scaled and matched for overall luminance using MATLAB’s Image Processing Toolbox.

**Figure 1.**
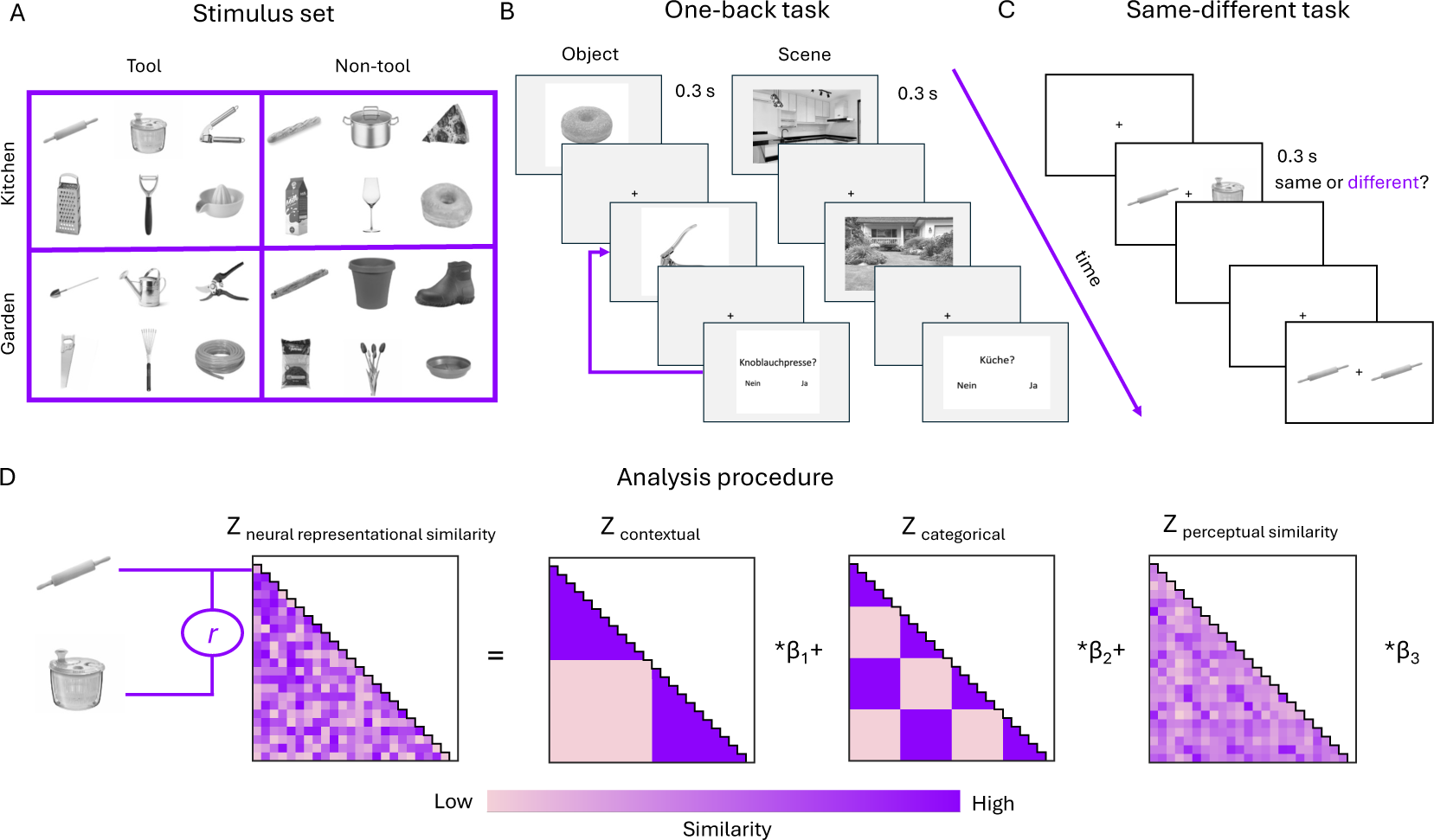
Stimuli, task, and schematic of analysis approach. (A) Stimulus set. 24 objects were used in the experiments. They stemmed from two contexts (kitchen and garden) and two categories (tool and non-tool). (B) Paradigm for both object (left) and scene (right) blocks. Participants saw isolated objects or scenes presented in a pseudo-random sequence and were asked to report whether an occasional word probe (in German) matched the previous image. (C) Same-different task procedure. Participants were asked to report whether two simultaneously presented stimuli were the same or different images. (D) Analysis approach. Neural RSMs constructed from the EEG and fMRI data were predicted from three model RSMs reflecting the objects’ similarities in contextual relations, (2) categorical relations, and (3) perceptual similarity. For each predictor, we estimated a beta weight in a linear model, separately for each participant. These beta weights indicated the contribution of contextual, categorical, and perceptual factors to temporally and spatially resolved neural activity.

We further assessed whether there were any visual differences between objects from the same and different contextual or categorical conditions. For this, we first computed Pearson correlations between layer-wise activation patterns in a VGG16 deep neural network (pre-trained on object categorization) for each pair of stimuli. This yielded a 24-by-24 representational similarity matrix (RSM) at every layer. Next, we created two predictor RSMs to model the image similarity matrix: (1) A contextual RSM, indicating whether object pairs belonged to the same context (e.g., both from the kitchen or garden) or different contexts; (2) A categorical RSM, indicating whether objects shared the same category (e.g., both tools or both non-tools) or different categories. We then vectorized all RSMs, retaining only the lower off-diagonal values. Next, the vectorized DNN RSM was predicted by a linear combination of the vectorized contextual and categorical RSMs in a regression model for each layer. Significant predictions were established from permutation tests, in which the rows and columns of the DNN RSM (1,000 iterations) were repeatedly shuffled to generate null distributions of beta estimates. The resulting p-values were corrected for comparisons across layers using false discovery rate (FDR) corrections. Within-condition (objects from the same context/category) correlations did not differ significantly from between-condition (objects from the different contexts/categories) correlations (largest contextual effect: β = −0.03, corrected *p* =.90, layer 4; largest categorical effect: β = 0.11, corrected *p* =.34, layer 15) effects, indicating no differential contribution of image features to the manipulated factors.

Furthermore, we used 24 scene images to test for shared representations between object images and their associated scene context. These images included 12 kitchen scenes and 12 garden scenes. None of our object stimuli featured in these scenes. The selected images were gray-scaled and matched for overall luminance using MATLAB’s Image Processing Toolbox. We then used a DNN to evaluate whether the objects shared a greater image similarity with the associated scene (e.g., rolling pin - kitchen) than the non-associated scene (e.g., rolling pin - garden). For this, we first computed Pearson correlations between layer-wise activation patterns in a VGG16 DNN, correlating activations for each object with activations for the two scenes (associated and non-associated). Next, we calculated the difference between the associated and non-associated conditions and tested the difference against zero using permutation t-tests with FDR correction (see above). Objects were equally correlated with the associated scene and the non-associated scenes (largest difference =.0089, corrected *p* =.769, layer 16), suggesting the objects did not systematically share image features with their associated scenes.

### Paradigm

In both EEG and fMRI experiments, participants first completed a questionnaire to assess their knowledge about the 24 selected objects. Here, the (German) words representing all objects were presented, and participants were asked whether they knew the object. If a participant did not know an object, the experimenter described it to ensure accurate identification during the task. During the main experiment, the participants first performed an object one-back task, indicating whether an occasionally presented word probe matched the previously shown object. Then, they performed the scene one-back task following the same logic (Fig. 1B). Finally, participants completed a co-occurrence questionnaire that covered all object pairs, designed to validate our assignment of objects to the two contexts. The questionnaire consisted of two parts: (1) object–scene co-occurrence and (2) object–object co-occurrence. In the first part, participants indicated where they typically encounter each object—either in a kitchen or a garden. In the second part, participants rated on a scale from 0 to 100 how frequently the two objects co-occur in daily life, and how similar their functions are— that is, whether the objects are used in a similar way and for a similar purpose. Across both experiments, all selected objects were rated as more frequently encountered within their assigned context, and co-occurrence between objects was consistently lower in the between-context condition compared to the within-context condition.

In the fMRI experiment, participants first completed an object practice block with 30 trials (24 images and 6 one-back trials) outside the scanner. Following an anatomical scan, we first ran an 8-minute functional localizer task comprising four conditions: fixation, object, scene, and scrambled objects. Each condition included 8 blocks, with 16 stimuli presented in each block. Stimuli were shown for 0.5 seconds each, followed by a 0.5-second interstimulus interval. The order of blocks was counterbalanced, and the same condition was never presented in consecutive blocks. After that, participants completed 10 object blocks and 2 scene blocks, each with 120 trials (96 images and 24 word-probes). Each object block began with a 6-second fixation dot and contained 96 images (featuring all object exemplars) with pseudo-randomization to prevent repetition of the same object (e.g., two rolling pin images shown consecutively) and 24 word-probe trials inserted at random positions in this sequence. In each scene block, we first pseudo-randomized 24 trials (featuring all scene exemplars) and subsequently inserted 6 word-probes (half kitchen and half garden); the procedure was applied four times, yielding 96 images and 24 probe trials for each block. For both types of blocks, each trial started with a 1.7-second fixation, followed by either a 0.3-second image or a 2.3-second word-probe display. The total fMRI session, including anatomical and localizer scans, lasted approximately 90 minutes, and the follow-up co-occurrence questionnaire again lasted around 30 minutes.

In the EEG experiment, participants sat in an illuminated room at a distance of 57 cm from the monitor. Participants first completed a practice object block with 30 trials. The following experiment consisted of 8 blocks of 240 trials. In each object block, 96 image trials were pseudo-randomized to prevent repetition of the same object, and 24 word-probes were inserted at random positions in this sequence. This procedure was applied two times, resulting in 192 image trials and 48 probe trials per block. After that, participants performed a scene practice block containing 12 scenes (half kitchen and half garden) and 4 one-back trials, followed by two experimental blocks of 240 trials each. In each scene block, we first pseudo-randomized 24 trials (featuring all scene exemplars) and subsequently inserted 6 word-probes (half kitchen and half garden); the procedure was applied eight times, yielding 192 images and 48 probe trials for each block. For both block types, each trial started with a fixation cross, which varied randomly between 0.8, 1, and 1.2 seconds, followed by a 0.3-second stimulus display. For the one-back trials, the word probes remained on screen until a response was made. Participants were instructed to maintain central fixation throughout, respond as quickly and accurately as possible in the one-back trials, and blink during ISIs. Participants could pause between blocks, and they could self-initiate the next block. The full EEG session lasted approximately 90 minutes, and the follow-up co-occurrence questionnaire lasted around 30 minutes.

### fMRI recording and preprocessing

MRI data was acquired using a 3T Siemens Magnetom PRISMA Scanner equipped with a 64-channel head coil. Functional images were obtained using T2*-weighted gradient-echo echo-planar imaging (EPI) with the following parameters: repetition time (TR) = 1850 ms, echo time (TE) = 30 ms, flip angle = 75°, in-plane voxel size = 2.2 mm x 2.2 mm x 2.2 mm, 58 slices with a 20% inter-slice gap (distance factor), field of view = 220 mm, matrix size = 100 × 100, and descending slice acquisition. Additionally, a high-resolution anatomical reference was collected using a T1-weighted MP-RAGE sequence with a voxel size of 1 mm^3^.

MRI data preprocessing was conducted using MATLAB and SPM12 (www.fil.ion.ucl.ac.uk/spm/). Functional images were first corrected for geometric distortions using the SPM FieldMap toolbox (Hutton et al., 2002) and subsequently realigned to account for head motion. Each participant’s structural T1-weighted image was coregistered to the mean of the realigned functional images. Normalization parameters for transformation to Montreal Neurological Institute (MNI) standard space—and their inverse—were then estimated.

For each run, functional data were modeled using a general linear model (GLM) that included separate regressors for the 24 objects. Additionally, six motion parameters obtained during realignment were included as nuisance regressors, resulting in a total of 30 regressors per run.

### fMRI regions of interest definition

fMRI analyses were focused on three regions of interest (ROIs): scene-selective parahippocampal place area (PPA), object-selective lateral occipital cortex (LOC), and early visual cortex (EVC). ROI masks were defined using individual localizer data constrained by group-level activation masks from functional brain atlases: For the PPA and LOC, we first combined left and right hemispheres and identified the top 500 voxels showing the highest t-values from the contrasts [scene > object + scrambled] and [object > scrambled], respectively, within regional masks derived from the atlas by Julian et al. (2012). For the EVC, all voxels from V1, V2, and V3 (including both dorsal and ventral subdivisions) were selected based on probabilistic maps from the Wang et al. (2015) atlas.

For analyses specifically targeting posterior and anterior PPA subregions, the PPA was further divided at MNI coordinate y = –42 following Baldassano et al. (2016). Within each subregion, the top 250 voxels were selected based on the scene > object + scrambled contrast from the localizer.

### fMRI RSA analysis

We used representational similarity analysis (RSA) to relate the contextual, categorical, and perceptual similarity of objects to their neural similarity (Fig. 1D). We first checked task performance to make sure participants followed the instructions. Task accuracy was 93.50% ± 1.12 % for object blocks and 93.23% ± 1.84 % for scene blocks, indicating that participants followed the instructions and paid attention to the objects and scenes. We averaged the fMRI beta maps from the first-level GLM analysis for each object across all exemplars and repeated trials across runs. Next, we computed the Pearson correlation of multi-voxel response patterns (i.e., patterns of beta values across voxels) between all pairs of objects in each ROI, resulting in a 24-by-24 RSM for each ROI. Next, we created three predictor RSMs to model the neural data: (1) A contextual RSM, indicating whether object pairs belonged to the same context (e.g., both from the kitchen or garden) or different contexts; (2) A categorical RSM, indicating whether objects shared the same category (e.g., both tools or both non-tools) or different categories; (3) a perceptual RSM, constructed from the reaction time of the same-different task (Fig. 1C). For this, an additional independent group of participants (N = 34) was recruited, indicating whether two simultaneously presented object images were the same or different. The RTs on the different-object condition were used as a measure of perceptual similarity, where longer RTs indicate higher perceptual similarity between the two objects (Jacob & Arun, 2020; Yeh & Peelen, 2022). The RSM was based on the average of the 34 participants. We then vectorized all RSMs, retaining only the lower off-diagonal values. Next, the vectorized neural RSMs were predicted by a linear combination of the vectorized contextual, categorical, and perceptual RSMs in a linear regression model (Fig. 1D). Finally, we tested whether the resulting beta estimates were larger than zero using one-sample t-tests, one-tailed. FDR correction was applied to control the multiple comparisons across the 9 tests (3 predictors and 3 ROIs).

To investigate the association between object and scene representations, we first averaged the fMRI beta values for each object and scene across all exemplars. Next, we correlated multi-voxel response patterns for each object with multi-voxel response patterns for the two scene categories using Pearson correlations. For each object, this resulted in one within-context (e.g., rolling pin and kitchen) and one between-context (e.g., rolling pin and garden) correlation. Next, we computed the difference between the within-context correlations and between-context correlations for each object and then averaged across objects. The resulting differences were tested against zero using one-sided t-tests at the group level.

### EEG recording and preprocessing

Electrophysiological signals were recorded using a 64-channel Easycap system with a Brain Products amplifier, sampled at 1,000 Hz. The setup included 56 electrodes distributed across both hemispheres (including FP1/FP2, AF3/AF4, AF7/AF8, F1/F2, F3/F4, F5/F6, F7/F8, FC1/FC2, FC3/FC4, FT5/FT6, FT7/FT8, FT9/FT10, C1/C2, C3/C4, C5/C6, T7/T8, CP1/CP2, CP3/CP4, CP5/CP6, TP7/TP8, P1/P2, P3/P4, P5/P6, P7/P8, PO3/PO4, PO7/PO8, PO9/PO10, and O1/O2) and seven electrodes along the midline (Fz, FCz, Cz, CPz, Pz, POz, and Oz). The ground electrode was placed at AFz, while Fz was used as the online reference. Electrodes FP1/FP2 and FT9/FT10 were repurposed for monitoring vertical and horizontal eye movements and were excluded from EEG analysis. Electrode impedances were maintained below 20 kΩ. Event triggers were transmitted from the stimulus presentation computer to the EEG acquisition system via a parallel port.

EEG data were preprocessed offline using the Fieldtrip toolbox (Oostenveld et al., 2011) in MATLAB (MathWorks). The TP10 channel, which exhibited excessive noise in most participants, was excluded and subsequently interpolated using the average signal from its neighboring electrodes. Ocular artifacts, including those from blinks and eye movements, were identified and removed through independent component analysis (ICA) combined with visual inspection of the components for each participant. The EEG data were then re-referenced to the average of all electrodes, excluding those related to ocular activity (FP1, FP2, FT9, and FT10), and segmented into epochs ranging from –100 ms to 500 ms relative to stimulus onset. Baseline correction was applied using the pre-stimulus interval (–100 to 0 ms). The data was then downsampled to 250 Hz for the following analysis. To focus on the object representations in the visual system, the anterior electrodes were excluded in the following analyses. All further analyses are based on the remaining 37 electrodes (Cz, CPz, Pz, POz, Oz, C1/C2, C3/C4, C5/C6, CP1/CP2, CP3/CP4, CP5/CP6, T7/T8, TP7/TP8, P1/P2, P3/P4, P5/P6, P7/P8, PO3/PO4, PO7/PO8, PO9/PO10, and O1/O2).

### EEG RSA analysis

To investigate how contextual, categorical, and perceptual factors influence the time course of object representations, we again used RSA, similar to the fMRI analysis (Fig. 1D). We first checked the task performance to make sure participants followed the instructions. Task accuracy was 89.92% ± 1.05 % for object blocks and 93.57% ± 0.92 % for scene blocks, indicating that participants followed the instructions and paid attention to the objects and scenes. For each time point in the EEG epochs, we first averaged the EEG signals for each object across all exemplars and trials. Next, we computed the Pearson correlation of EEG signals between all pairs of objects at each time point. This yielded a 24-by-24 neural representational similarity matrix (RSM) at every time point (250Hz resolution). Next, we created three predictor RSMs to model the neural data: (1) A contextual RSM, indicating whether object pairs belonged to the same context (e.g., both from the kitchen or garden) or different contexts, (2) a categorical RSM, indicating whether objects shared the same category (e.g., both tools or both non-tools) or different categories, and (3) A perceptual RSM, constructed from the reaction time of the same-different task. (in the same way as for the fMRI analysis). We then vectorized all RSMs, retaining only the lower off-diagonal values. Next, vectorized neural RSMs were predicted by a linear combination of the vectorized contextual, categorical, and perceptual RSMs in a linear regression model. Finally, we tested whether the resulting beta estimates were greater than zero (one-sided) at each 4-ms timepoint within the 0–500 ms window following stimulus onset, using cluster-based non-parametric permutation t-tests (Maris & Oostenveld, 2007). Specifically, t-tests were first performed at each time point. Time points exceeding a predefined threshold (*p* <.05) and occurring consecutively were grouped into clusters. For each cluster, the sum of the t-values was computed and compared against a null distribution generated from 1,000 Monte Carlo random permutations. Statistical significance was determined using a cluster-level correction for multiple comparisons (one-tailed, p <.05).

To investigate the association between object and scene representations, we first computed event-related potentials (ERPs) for each object and scene across all exemplars. Next, we correlated ERP patterns (across electrodes) for each object with ERP patterns for the two scene contexts using Pearson correlations, separately for each 4-ms time point. For each object and time point, this resulted in one within-context (e.g., rolling pin and kitchen) and one between-context (e.g., rolling pin and garden) correlation. Next, we computed the difference between the within-context correlations and between-context correlations for each object and then averaged across objects. The resulting differences were tested against zero using one-sided t-tests at the group level with non-parametric cluster correction (see above).

## Results

Here, we applied RSA to fMRI and EEG data to investigate where and when contextual associations shape object representations, while controlling for perceptual and categorical similarity. We first focus on the fMRI results, revealing which brain regions represent contextual object relations.

### Contextual object relations are represented in object- and scene-selective cortex

By performing RSA on the fMRI data, we aimed to investigate which visual cortex regions reflect contextual relations, focusing on PPA, LOC, and EVC (Fig. 2A). We observed contextual effects in PPA [*t*(31) = 2.14, FDR-corrected *p* =.036] and LOC [*t*(31) = 2.38, FDR-corrected *p* =.027], suggesting a representation of contextual object information in both object- and scene-selective cortex. No contextual effects were observed in EVC [*t*(31) = 0.23, FDR-corrected *p =*.460]. Categorical effects were only observed in LOC [*t*(31) = 6.07, FDR-corrected *p* <.001] but not in PPA[*t*(31) = 1.00, FDR-corrected *p =*.243] and EVC [*t*(31) = −1.21, FDR-corrected *p =*.883]. Finally, perceptual effects were observed in LOC [*t*(31) = 5.58, FDR-corrected *p* <.001] and EVC [*t*(31) = 3.85, FDR-corrected *p* <.001], but not in PPA [*t*(31) = −1.73, FDR-corrected *p =*.953]. Together, these results reveal an emergence of both contextual and categorical effects in object processing regions, contrasting with only contextual information represented in scene-selective PPA.

**Figure 2.**
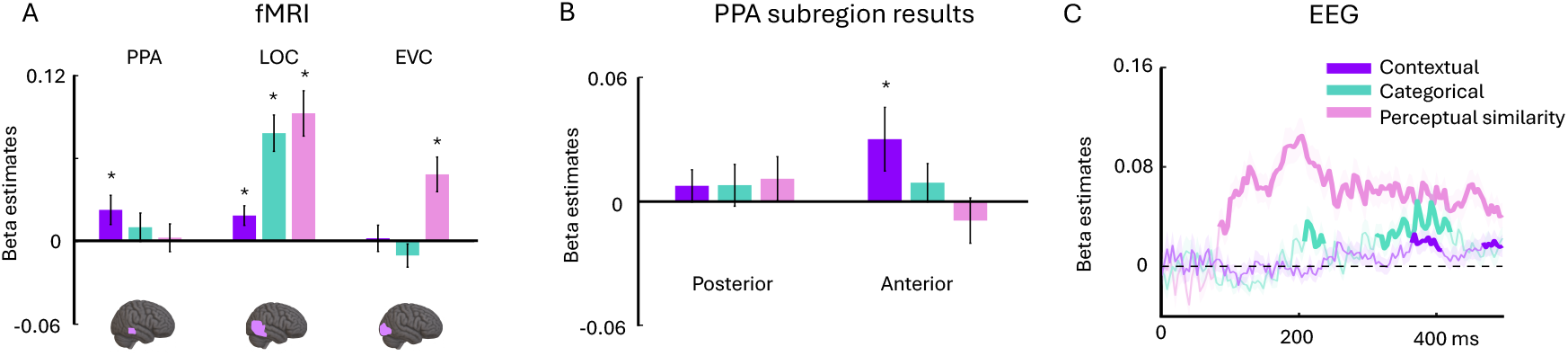
RSA results. (A) fMRI results. Contextual effects were observed in PPA and LOC, categorical effects in LOC, and perceptual effects in LOC and EVC. * indicates FDR-corrected *p* < 0.05, one-tailed. (B) fMRI results in PPA subregions. Contextual effects were only found in the anterior PPA. * indicates *p* < 0.05, one-tailed. (C) EEG results. Contextual effects emerged at 364 ms and again at 468 ms after stimulus onset, later than the perceptual effects (from 84 ms) and categorical effects (from 208 ms). Bold lines indicate statistical significance (*p* < 0.05, cluster-level, one-tailed).

Next, we examined how contextual effects emerge across PPA subregions. Prior studies have shown that the posterior PPA is more responsive to visually driven information, whereas the anterior PPA is associated with conceptual processing (Baldassano et al., 2013, 2016). Following Baldassano et al. (2016), we divided the PPA at MNI coordinate y = –42 and applied the identical RSA to each subregion (see Methods fMRI RSA). Interestingly, we found a significant contextual effect only in the anterior PPA [*t*(31) = 1.95 *p* =.030], but not in the posterior PPA [*t*(31) = 0.96, *p* =.173], and the contextual effect was larger in anterior PPA than in posterior PPA [*t*(31) = 2.01, *p* =.052, two-tailed] (Fig. 2B).

These results indicate that contextual relations are represented in object-selective LOC and scene-selective PPA, but not in EVC. Together with previous fMRI studies (Bar & Aminoff, 2003; Bonner & Epstein, 2021), this pattern suggests that representations of contextual object relations arise from relatively late interactions among representations across high-level visual cortex. To characterize the precise timing of contextual relation representations, we next performed RSA on the EEG data.

### Contextual object relations are represented during the late stages of perceptual processing

We investigated the time course of contextual object representations using RSA on the EEG data. Contextual effects emerged in two temporal clusters. First, from 364 ms to 411 ms after stimulus onset (cluster *p*=.008), and second, from 464 ms after stimulus onset to the end of the epoch (cluster *p*=.040; Fig. 2C). Critically, these contextual effects emerged later than perceptual effects, which were evident from 84 ms after stimulus onset (cluster *p* <.001), and later than categorical effects, which were evident from 208 ms (cluster *p* =.044) and from 312 ms (cluster *p* <.001) after stimulus onset. These temporal dynamics reveal that, upon viewing an object, there is a sequential emergence of information, with perceptual information followed by categorical information (here: whether the object is a tool or not), and finally contextual information (here: whether an object is found in the kitchen or garden).

### Representations of contextual object relations may partly reflect the activation of scene representations

Next, we examined whether the isolated objects and scenes evoke shared representations, which would indicate that contextual representations are mediated by an automatic co-activation of associated scene representations. Using RSA on the fMRI and EEG data, we identified the spatiotemporal signatures of this putative co-activation. Specifically, we compared response patterns between objects and their associated scenes (within-context correlation) versus the non-associated scenes (between-context correlation) (Fig. 3A).

**Figure 3.**
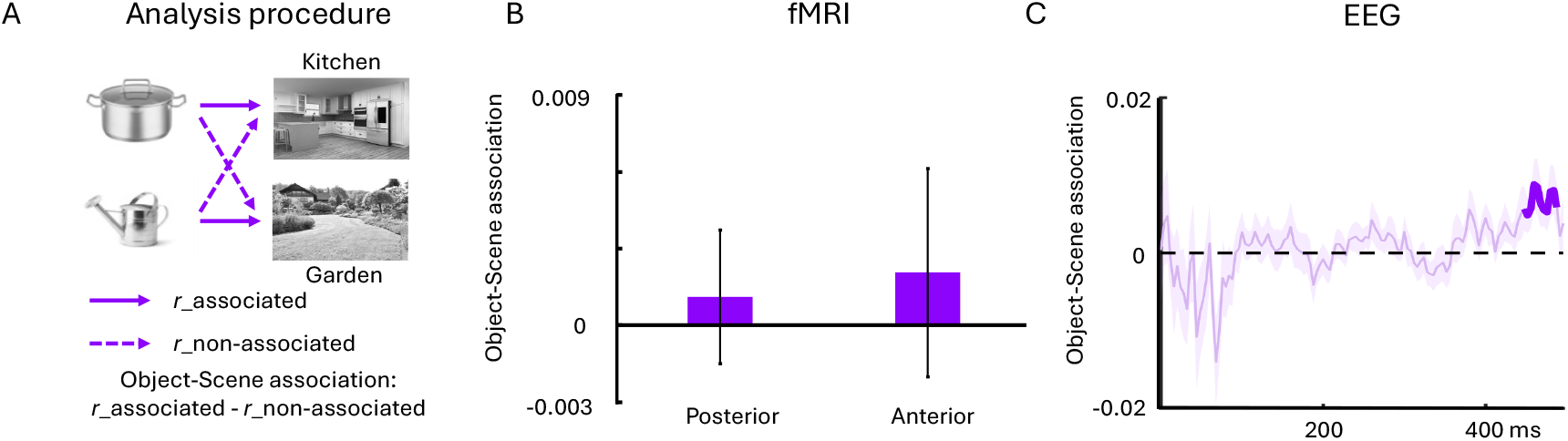
Results from the scene co-activation RSA. (A) Schematic of the analysis approach. We first computed correlations between each object and its associated scene (within-context correlation) and the non-associated scene (between-context correlation). Subtracting these two correlations yielded a measure of category-specific object-scene association during object viewing. (B) fMRI results. No significant object–scene associations were observed in the anterior and posterior PPA subregions. (C) EEG results. A significant cluster of scene association emerged around 450 ms after stimulus onset (cluster *p* < 0.05).

For the fMRI analysis, we focused on the posterior and anterior PPA subdivisions, given that representations of scene categories should form in the PPA (Walther et al., 2009) and given that the previous contextual relation effects emerged in the anterior portion of the PPA. However, we did not observe significant effects, neither in the anterior PPA [*t*(31) = 0.51, corrected p =.308] nor in the posterior PPA [*t*(31) = 0.42, *p* =.338] (Fig. 3B). Further, no effects emerged in regions outside of the PPA (EVC, LOC, all *t*(31) < 1.31, *p* >.1).

Performing this analysis on the EEG data, we found significantly higher within-context than between-context correlations from 452 ms after stimulus onset (cluster *p* =.011; Fig. 3C). This suggests that late representations of contextual object relations (i.e., the second cluster found in the previous analysis; Fig. 2C) are partly related to the co-activation of associated scene representations.

In sum, our EEG results suggest that late representations of contextual object relations (emerging around 460 ms after stimulus onset) may partly relate to the co-activation of associated scenes. However, we were not able to localize such representations in our fMRI analysis. Further studies are needed to fully clarify whether and how the co-activation of scene representations contributes to contextual codes in object vision.

## Discussion

In this study, we used fMRI and EEG to investigate where and when contextual associations shape neural representations of real-world objects. Applying RSA on fMRI and EEG signals, we found that contextual information is represented in object-selective areas LOC and scene-selective anterior PPA. Such representations emerged during late stages of visual object processing, from around 360 ms after stimulus onset. Contextual representations first reflected similarities among objects from the same scene context, and subsequently showed signs of the co-activation of object representations and their associated scene category representations.

The finding that contextual relations are encoded in scene-selective PPA, is consistent with the fMRI results of Bonner and Epstein (2021). By using a tightly controlled stimulus set and systematically accounting for categorical and perceptual similarity in our analysis, our results demonstrate a genuine representation of contextual associations in PPA, rather than shared perceptual or categorical features. We also found contextual object representations in LOC, consistent with Bar and Aminoff’s (2003) report of objects with stronger contextual associations also activating LOC more strongly. The contextual effects in LOC may reflect the co-activation of object concepts through learned object-to-object relations encountered in real-world environments (Kim & Biederman, 2011; Kaiser et al., 2019; Vo, 2021).

Within the PPA, we found that contextual relations were encoded in the anterior but not the posterior PPA. While the posterior PPA is more strongly connected to occipital areas, supporting the processing of visual features such as spatial layout (Baldassano et al., 2013, 2016; Nasr et al., 2014; Lescroart & Gallant, 2019), the anterior PPA is more strongly connected to the hippocampus, particularly its anterior part, which is involved in episodic memory encoding and retrieval, environmental representation, and scene construction for recall and imagination (Baldassano et al., 2016; Zeidman & Maguire, 2016). The emergence of contextual representations in anterior PPA suggests that the brain retrieves relations among co-occurring objects from episodic memory, likely to mentally construct associated scenes and episodes, which may in turn facilitate the recognition of isolated objects (Bar & Aminoff, 2003; Bar, 2004; Hayes et al., 2007; Bar et al., 2008; Aminoff et al., 2013; Steel et al., 2023). On this view, contextual representations in PPA may reflect more abstract relational processing rather than the reactivation of concrete visual features such as spatial scene layouts (Epstein, 2008; Mullally & Maguire, 2011, 2013).

Our EEG results revealed a sequential emergence of three effects: perceptual effects appeared first (from 84 ms post-stimulus), followed by categorical effects (from 208 ms), and finally contextual effects, which emerged later in processing (from 364 ms post-stimulus). We observed two distinct temporal clusters during which contextual information was represented: between 364 ~ 411 ms and between 464 ms to the end of the epoch. The sequence of emergence aligns with the hierarchical organization of the visual system, in which early visual cortices process low-level features, while higher-order regions encode more semantic information (Peelen & Caramazza, 2012; Peelen et al, 2013; Clarke et al., 2013; Cichy et al., 2014; Proklova et al., 2016, 2019; Thorat et al., 2019). During this semantic coding stage, contextual effects emerged later than categorical effects, suggesting that contextual processing requires prior computation of object categories, rather than reflecting a form of rapid visual priming among co-occurring objects. The delayed contextual effect is also consistent with MEG findings showing that high-level scene representations (Cichy et al., 2017) and scene-based contextual facilitation of object recognition (Brandman & Peelen, 2017) arise relatively late in visual processing. This indicates that inferring scene context requires additional processing time, perhaps necessitated by memory retrieval from higher-order cortical regions. Here, further studies are needed to clarify the role of memory processes in the formation of contextual relation representations.

It is worth noting that two recent EEG studies produced seemingly inconsistent results with ours. First, Kim et al. (2025) did not observe more similar neural representations between objects that are contextually related, here measured by the frequency of object co-occurrence. However, not all stimuli in their study were strongly tied to a specific scene context (e.g., toys, clothing, or body parts can appear in many contexts), and associations to a common scene context may be needed to produce reliable effects. Second, Kallmayer et al. (2024) reported shared representations for objects from the same scene context (e.g., toothbrush and sink) during a much earlier time window (128 – 164 ms post-stimulus) than we did. It is possible that these earlier effects are driven by categorical factors, as categorical similarity was not accounted for in that study.

The second temporal cluster during which contextual information was represented (from 464 ms to the end of the epoch) overlapped with the time window during which we observed shared representations between objects and their associated scenes (from 452 ms to the end of the epoch). This suggests that there might be two distinct stages of contextual representation: An initial co-activation among contextually related objects may be followed by a co-activation of associated object and scene representations. However, we found no evidence for such a co-activation between objects and scenes in the fMRI, neither in LO and PPA nor in a spatially unconstrained searchlight analysis. Previous studies found no shared representations between scenes and individual objects in the PPA either (MacEvoy & Epstein, 2011; Kaiser & Peelen, 2018), but findings in LOC are mixed: Kaiser and Peelen (2018) reporting no shared representations, while MacEvoy and Epstein (2011) could successfully cross-decode between objects and their associated scenes. Here, it is important to note that the MacEvoy and Epstein (2011) study featured the same objects in the individual object and scene conditions, while our study (as well as Kaiser & Peelen, 2018) used scenes that did not contain the individual objects. While the absence of scene-object associations in the fMRI is therefore in line with previous fMRI findings, the difference between our EEG and fMRI experiments is harder to reconcile. One possibility is that EEG signals more faithfully capture activity across distributed regions and thus unveil the underlying interactions between object- and scene-processing regions that drive object-scene associations. At this point, further studies are needed to fully understand whether such co-activations between objects and scenes are reliable and, if so, where they are localized in the brain.

Previous behavioral studies demonstrated that context frames facilitate real-world object recognition (Oliva & Torralba, 2007). Such effects manifest as facilitation effects among objects, where target objects are recognized more accurately when presented with context-consistent non-target objects (Auckland et al., 2007), as well as among objects and scenes, where consistent scene context enhances the identification of objects, and vice versa (Davenport & Potter, 2004; Brandman & Peelen, 2017; Leroy et al., 2020; Chen et al., 2022). Our current results suggest that such contextual facilitation is supported by the co-activation of contextually associated object representations and shared scene category representations in visual cortex. These co-activations can sharpen neural representations of ambiguous objects (Brandman & Peelen, 2017; Quek et al., 2025) and scenes (Brandman & Peelen, 2023) and lower the perceptual threshold for upcoming stimuli by pre-activating representations of associated objects and scenes (Tang et al., 2022).

## Conclusion

In summary, our findings show that contextual associations of objects are represented in object- and scene-selective regions of the human ventral visual system and emerge during late stages of object processing. Such contextual relation representations emerged for isolated objects, and without a task that encourages their formation, suggesting that objects automatically activate context frames that support visual cognition in real-world environments.

## Acknowledgements

The authors thank Sirine Nouira for part of the EEG data collection. MR-imaging for this study was performed at the Bender Institute of Neuroimaging (BION) at the Justus Liebig University Giessen, Germany.

## Notes

### Data and materials availability

All study materials supporting this research are publicly available: https://osf.io/t62s3/files/osfstorage

